# Comparative analysis of coronavirus genomic RNA structure reveals conservation in SARS-like coronaviruses

**DOI:** 10.1101/2020.06.15.153197

**Authors:** Wes Sanders, Ethan J. Fritch, Emily A. Madden, Rachel L. Graham, Heather A. Vincent, Mark T. Heise, Ralph S. Baric, Nathaniel J. Moorman

**Affiliations:** University of North Carolina at Chapel Hill, Department of Microbiology and Immunology, NC, USA; University of North Carolina at Chapel Hill, Lineberger Comprehensive Cancer Center, NC, USA; University of North Carolina at Chapel Hill, School of Public Health, NC; University of North Carolina at Chapel Hill, Department of Genetics, NC, USA, USA

**Author notes:** Corresponding author Correspondence and requests for materials should be addressed to Nathaniel J. Moorman. Duke University, Department of Surgery, NC, USA.

## Abstract

Coronaviruses, including SARS-CoV-2 the etiological agent of COVID-19 disease, have caused multiple epidemic and pandemic outbreaks in the past 20 years^1–3^. With no vaccines, and only recently developed antiviral therapeutics, we are ill equipped to handle coronavirus outbreaks^4^. A better understanding of the molecular mechanisms that regulate coronavirus replication and pathogenesis is needed to guide the development of new antiviral therapeutics and vaccines. RNA secondary structures play critical roles in multiple aspects of coronavirus replication, but the extent and conservation of RNA secondary structure across coronavirus genomes is unknown^5^. Here, we define highly structured RNA regions throughout the MERS-CoV, SARS-CoV, and SARS-CoV-2 genomes. We find that highly stable RNA structures are pervasive throughout coronavirus genomes, and are conserved between the SARS-like CoV. Our data suggests that selective pressure helps preserve RNA secondary structure in coronavirus genomes, suggesting that these structures may play important roles in virus replication and pathogenesis. Thus, disruption of conserved RNA secondary structures could be a novel strategy for the generation of attenuated SARS-CoV-2 vaccines for use against the current COVID-19 pandemic.

## Main

Severe acute respiratory syndrome coronavirus (SARS-CoV), Middle Eastern respiratory syndrome coronavirus (MERS-CoV), and SARS-CoV-2, the etiological agent of the current COVID-19 pandemic, have caused widespread disease, death, and economic hardship in the past 20 years^1^, highlighting the pandemic potential of the CoV genus. While recently developed antivirals show promise against MERS and SARS-CoV-2, further understanding of coronavirus molecular virology is necessary to inform the design of more effective antiviral therapeutics and vaccines^6, 7^.

RNA structures in the ~30kb of single-stranded RNA^8^ genomes of Coronavirus play important roles in coronavirus replication^9, 5, 10, 11, 12, 13^. Given the length of coronavirus RNA genomes, additional RNA structures likely exist that regulate CoV replication and disease^14^. In this study, we used selective 2’-hydroxyl acylation by primer extension and mutational profiling (SHAPE-MaP)^15^ to identify highly stable, structured regions of the SARS-CoV, MERS-CoV, and SARS-CoV-2 genomes. Our results revealed novel areas of RNA structure across the genomes of all three viruses. SHAPE-MaP analysis confirmed previously described structures, and also revealed that SARS-like coronaviruses contain a greater number of highly structured RNA regions than MERS-CoV. Comparing nucleotide variation across multiple strains of each virus, we find that highly variable nucleotides rarely impact RNA secondary structure, suggesting the existence of selective pressure against RNA secondary structure disruption. We also identified dozens of conserved highly stable structured regions in SARS-CoV and SARS-CoV-2 that share similar structures, highlighting the possible importance of these stable RNA structures in virus replication.

### Stable RNA secondary structure is pervasive throughout coronavirus genomes

To identify regions of significant RNA secondary structure in the genomes of MERS-CoV, SARS-CoV, and SARS-CoV-2 we performed selective 2′-hydroxyl acylation analyzed by primer extension and mutational profiling (SHAPE-MaP) analysis of virion-associated RNA^15, 16^ (**Fig 1A-C; Supplemental Table 1**). Highly stable structured regions likely to maintain a single confirmation were identified using high confidence cutoffs for SHAPE reactivity (<0.3, blue bars) and of Shannon entropy (<0.04, black bars). We identified 85 highly stable regions in the MERS-CoV genome (**Fig 1A**), 123 stable regions in the SARS-CoV genome (**Fig 1B**), and 139 stable regions within the SARS-CoV-2 genome (**Fig 1C**). Stable structured regions were present throughout the coding and non-coding regions of each genome. SARS-CoV and SARS-CoV-2 exhibited greater RNA structuredness across their genomes compared to MERS-CoV. This was not due to differences in overall GC content or bias in average SHAPE reactivity (**Supplemental Fig 1A**). Thus, SARS-like coronavirus genomes contain greater overall stable RNA structuredness than the MERS-CoV genome.

**Figure 1.**
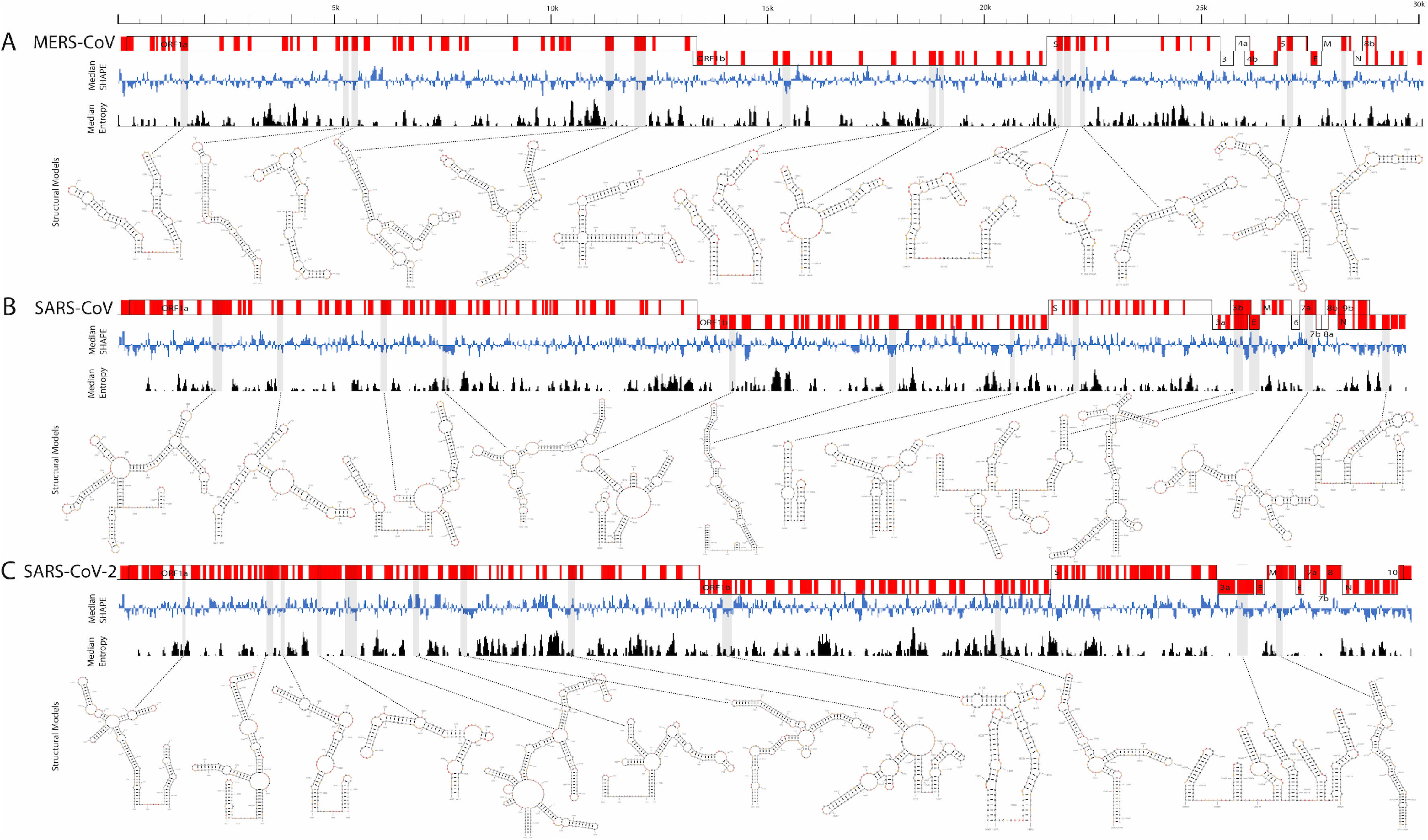
RNA secondary structure is pervasive throughout coronavirus genomes. Schematic of genome architecture for MERS-CoV (**A**), SARS-CoV (**B**), and SARS-CoV-2 (**C**). Local median SHAPE reactivity (55-nt window) compared to the global median reactivity is shown in blue. Median Shannon entropy (55-nt window) compared to the global entropy is represented in black. Areas of significantly high RNA secondary structuredness (merged SHAPE reactivity and Shannon entropy data, see Methods) are highlighted in red for each genome. Examples of highly stable, RNA secondary structures are shown for each virus. Grey bars below the genome schematics denote these structures are located.

We used the SHAPE reactivity to create a structural model for stable RNA elements in each genome (**Fig 1A-C; Supplemental Figure 5, Supplemental Table 2**). The top twenty most stable structures for each virus were present in the coding region, and largely consisted of hairpin and bulged stem loop structures. The most highly stable RNA structures were present in the regions encoding open reading frame (ORF) ORF1a, ORF1b and Spike. The majority of the most highly stable structures for all coronaviruses were present within ORF1a, while the number of highly stable structures in ORF1b ranged from three in SARS-CoV to eight in MERS-CoV (**Supplemental Fig 1B**). All coronaviruses had multiple highly stable structures in the Spike ORF. These data highlight the pervasive presence of highly stable RNA structure throughout coronavirus genomes.

### SHAPE-Map RNA structure prediction recapitulates known coronavirus structural elements

The RNA structural elements within the 5’ and 3’ UTRs of the related murine hepatitis coronavirus (MHV) genome have been extensively characterized ^5, 17,18^. Our SHAPE-MaP analysis recapitulated previously described 5’ and 3’ UTR RNA secondary structures. Within the 5’ UTR of MERS-CoV, SARS-CoV and SARS-CoV-2, we identified the conserved stem loop (SL) SL1, SL2, SL4 and SL5ABC RNA elements (**Fig 2A**). Consistent with previous modeling^5^, both SARS-like coronaviruses contained SL3 upstream of SL4, while this hairpin was absent from MERS-CoV. A similar pattern was observed for structures within the 3’UTR (**Fig 2B**). Each virus exhibited a bulged stem loop (BSL) structure, followed by a pseudoknot (PK) and each genome ends in a long hypervariable (HVR) bulged stem loop structure, confirming the conservation of these RNA secondary structures across coronaviruses.

**Figure 2.**
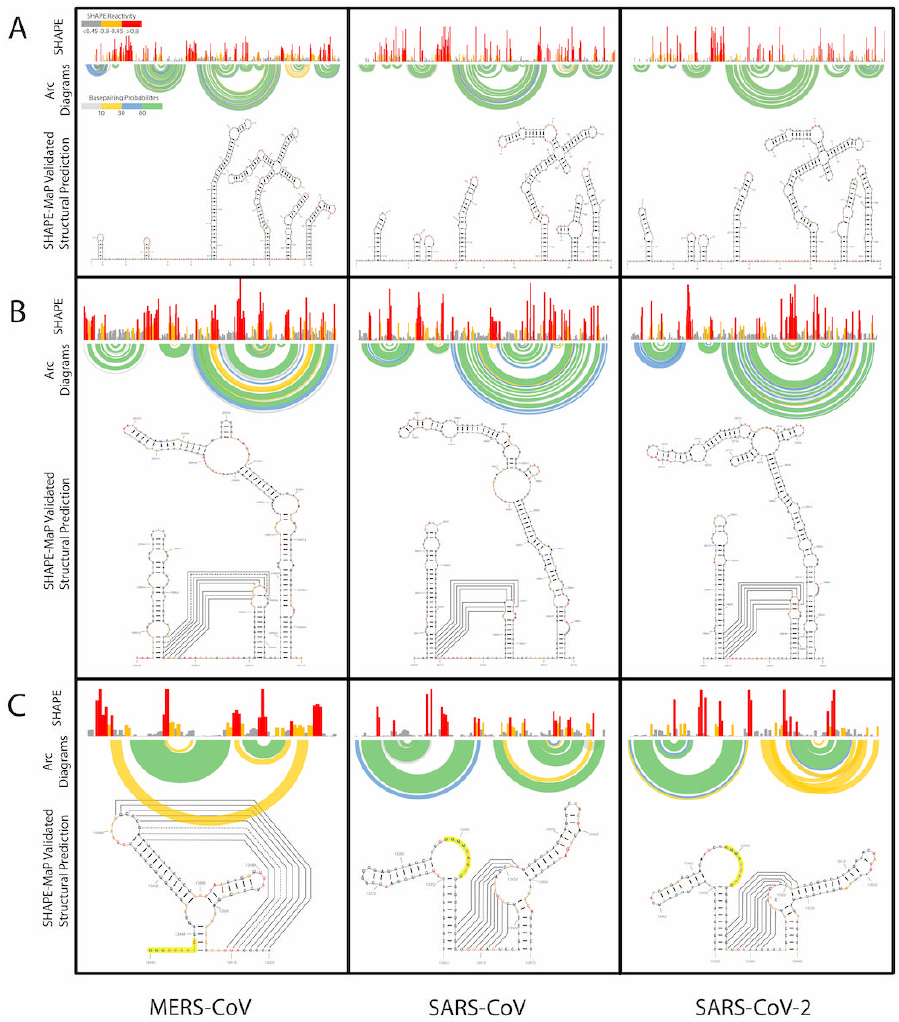
Coronaviruses exhibit conserved secondary structure in functional RNA elements. SHAPE reactivity (top panel), base pair probability arc diagrams (middle panel), and predicted RNA secondary structure models for the 5’ UTR (**A**), 3’ UTR (**B**), and structures surrounding the programmed −1 ribosomal frameshift site (**C**) for MERS-CoV, SARS-CoV, and SARS-CoV-2 are shown. Black connected lines represent previously described pseudoknot tertiary interactions. The ‘slippery sequence’ that promotes ribosomal pausing is highlighted in yellow.

Coronaviruses utilize a programmed −1 ribosomal frameshift to produce the ORF1a/1b fusion protein. A stable RNA pseudoknot upstream of the frameshift position is necessary for the frameshift to occur^12, 13^, and our data supports the presence this pseudoknot in the same position in all three coronaviruses (**Fig 2C**). Interestingly, the MERS-CoV two-stemmed pseudoknot is distinct from the single-stemmed pseudoknot present in both SARS-like coronaviruses. Together, these results demonstrate the capability of SHAPE-MaP analysis to accurately identify RNA structures in coronavirus genomic RNA.

Transcriptional regulatory sequences (TRSs) are conserved, short sequences (6-8 nucleotides) within coronavirus genomes that act as cis-regulatory elements of transcription^18, 19, 20^. While RNA secondary structure has been suggested plays a role in TRS regulation of transcription, most TRS elements were not associated with areas of high RNA structuredness (**Supplemental Fig 2A-D**). In fact, only a single TRS from MERS-CoV and SARS-CoV were associated with highly structured RNA regions, while four TRSs from SARS-CoV-2 were associated. Thus, TRS elements are not consistently associated with highly stable RNA structures, suggesting that RNA secondary structure may not play a role in TRS recognition.

### Covariance and MUSCLE alignment analysis reveal selective pressure for coronavirus RNA secondary structure

To determine if RNA structuredness is a driver of mutational variance across virus strains, we performed multiple sequence comparison by log-expectation (MUSCLE) alignment analysis of 350 MERS-CoV, 141 SARS-CoV, and 1,542 SARS-CoV-2 genomes (**Supplemental Table 3**). As a whole, highly structured regions within SARS-CoV exhibited significantly less nucleotide variance than areas of lower structuredness, suggesting these regions may be under higher selective pressure. There was no significant difference in nucleotide variance within MERS-CoV or SARS-CoV-2 genomes, however 72.9% of highly structured regions in MERS-CoV and 87.8% of highly structured regions in SARS-CoV-2 had lower variance than regions of lower structuredness. Eighteen nucleotides in MERS-CoV, 3 nucleotides in SARS-CoV, and 8 nucleotides in SARS-CoV-2 showed significant variance across all strains and were located in structured regions (**Supplemental Table 4**). Of these variants, 72% in MERS-CoV, 100% in SARS-CoV, and 75% in SARS-CoV-2 occur in unpaired nucleotides in the stable RNA secondary structure, suggesting that the majority of polymorphisms in highly structured regions across coronavirus strains maintain RNA secondary structures.

We performed covariation analysis to further assess conservation of structuredness. This analysis determines if two nucleotide changes occurred in tandem to conserve RNA secondary structure. 21 structured regions in MERS-CoV strains, 1 structured region in SARS-CoV strains, and 4 structured regions in SARS-CoV-2 strains contained significant covarying nucleotide changes that conserved RNA secondary structure (**Supplemental Table 5**). Together, these results suggest that regions of RNA structuredness and specific RNA secondary structures are conserved within each virus family, and these structured regions may be under selective pressure

### SARS-CoV-2 and SARS-CoV structured regions are highly conserved

We next sought to identify conserved, highly stable RNA structures across all three viruses. Surprisingly, outside of the previously described conserved 5’ and 3’ UTRs, we identified only a single conserved structure across all three genomes, present within the RNA region encoding nsp16 (**Supplemental Fig 3**). However, we found 98 areas of overlapping highly structured regions when comparing only the SARS-CoV and SARS-CoV-2 genomes (**Fig 3A & Table 7**). Within conserved regions of structuredness in the coding region of the SARS-CoV-2 genome we identified 65 regions that contained similar RNA secondary structures (**Fig 3B**). The majority of these structures (26 total) were within ORF1b, in contrast to the distribution of the most stable structures within a respective genome (**Supplemental Fig 4**).

**Figure 3.**
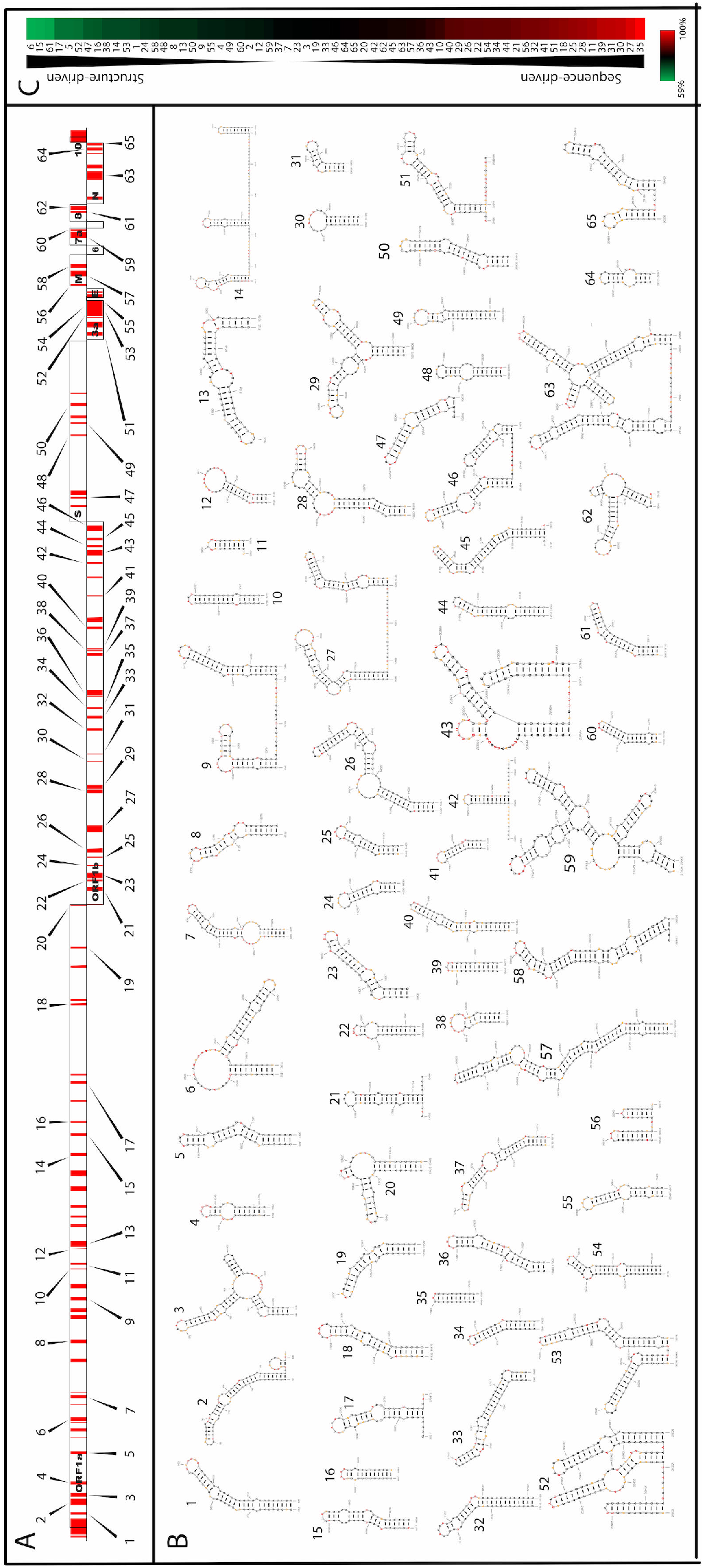
SARS-like CoV contain highly conserved regions of RNA structuredness. (**A**) SARS-CoV-2 genomic architecture schematic. Conserved regions of RNA structuredness between SARS-CoV-2 and SARS-CoV are highlighted in red. Conserved highly stable RNA secondary structures are denoted with arrows. (**B**) Representative structural models conserved, highly stable RNA structures from (A) are shown. (**C**) The percent nucleotide conservation between SARS-CoV and SARS-CoV-2 for each highly stable conserved RNA structure is shown in a heat map. The highest conservation (100%) is shown in red, average nucleotide conservation (79%) is represented in black, and the lowest conservation (59%) is shown in green.

SARS-CoV and SARS-CoV-2 share a 79% sequence homology, thus, conserved RNA structures may be driven by sequence homology rather than conservation of RNA secondary structure (**Supplemental Table 6**). To determine which RNA structures were likely conserved based on RNA secondary structure rather than sequence homology we calculated the average percent nucleotide conservation between SARS-CoV and SARS-CoV-2 for each conserved structure and compared this to the average nucleotide conservation of the two genomes (**Fig 3C**). Similar secondary structures that exhibit greater than average nucleotide homology are likely driven by sequence homology, while similar structures that show lower than average nucleotide homology are likely conserved based on preservation of RNA secondary structure. We found 18 total structures (27.7%) with lower than average nucleotide homology, with structure 6 showing the lowest sequence homology (59.4%). These data suggest that nearly one third of conserved similar RNA secondary structures between SARS-CoV and SARS-CoV-2 are conserved based on RNA secondary structure.

Lastly, we assessed nucleotide covariation within conserved SARS-like CoV RNA secondary structures. SARS-CoV-2 genomes were compared to MERS-CoV and SARS-CoV strains and covarying nucleotides that support SARS-CoV-2 structures were identified (**Supplemental Table 5**). In line with a lack of structure conservation, no MERS-CoV strains showed covarying nucleotides that supported SARS-CoV-2 structures. In contrast, we identified 10 significant covariation events within SARS-CoV genomes that supported SARS-CoV-2 structures, two of which occurred within the viral UTRs. Five covariation events were observed in non-conserved regions of structuredness, while three covariation events occurred within conserved regions of structuredness. These results show that conservation of specific RNA structures may extend outside of conserved regions of structuredness, and overall highlight the high degree of structural conservation between SARS-CoV and SARS-CoV-2.

## Discussion

Using SHAPE-MaP analysis we found that members of the *Coronaviradae*, MERS-CoV, SARS-CoV, and SARS-CoV-2, contain highly stable, structured RNA regions throughout their genomes. SARS-like coronavirus genomes were more highly structured than the MERS-CoV genome, suggesting that even within the betacoronavirus family, RNA structuredness may be unique to viral species. Importantly, SHAPE-MaP analysis recapitulated previously described RNA structures^5^, providing confidence that SHAPE-MaP identifies bona fide RNA secondary structures. Outside of structurally conserved noncoding regions of the genome, only a single RNA structure was conserved across all three viruses. However, 65 conserved highly stable structured regions were found that contained similar RNA structures when comparing SARS-CoV and SARS-CoV-2 (**Supplementary Fig 3**). While some structures were conserved based on sequence homology alone, the conservation in nearly a third of these structures appears to be driven by structural conservation. Alignment and covariance analysis of >1000 SARS-CoV and SARS-CoV-2 strains revealed greater levels of sequence conservation in SARS-like CoV structured RNA regions, and maintenance of RNA secondary structures in highly stable structures. This suggests these regions are under greater selective pressure than their non-structured counterparts, and their likely importance in SARS-like coronavirus replication.

In contrast, we found distinct structural differences between MERS-CoV and the SARS-like CoV, even within the known functional RNA structures (e.g. 5’ and 3’UTR, ribosomal frameshift element; **Fig 2A, 2C**). Such subtle, but distinct, changes in RNA structure could drive phenotypic differences in replication of distinct virus species, even when those structures are within conserved regions of RNA structure.

We also found significant conservation of both structuredness and RNA secondary structures within the SARS-like CoV (**Fig 3A,B**). While much of this conservation was due to sequence homology between the two viruses, we found that almost a third of conserved structures show lower than average sequence homology as compared to the rest of the genome, suggesting a functional role for RNA secondary structure (**Fig 3C**). Further, in structures that did show high sequence homology, we found evidence of covariance conservation of RNA structures, suggesting sequence variation constraints within the context of an RNA structure. In conjunction with the fact that structured regions in SARS-CoV showed lower sequence variance than non-structured regions, we hypothesize that RNA structuredness and specific RNA secondary structures exert a selective pressure on sequence diversity. This would suggest that these structures play important roles in SARS-like CoV replication, and possibly pathogenesis. This suggests that disrupting these such conserved structures could be a promising strategy for the development of live-attenuated vaccines. Understanding the mechanism by which these novel RNA structures regulate SARS-CoV-2 replication could lead to new antiviral therapeutic targets and inform rational vaccine design.

## Supplemental Files and Figures

Supplemental Figure S1-S5

Supplemental Tables T1-T7

## Methods

### Virus growth and purification

MERS-CoV Jordan-N3/2012 isolate MG167 (accession #KJ614529), was gown in Vero-81 cells. Vero-81 cells were cultured to ~80% confluence in T175 flasks. Immediately prior to infection, the culture medium was aspirated and replaced with Opti-MEM with 4% Hyclone FetalClone II (Cytiva). Cells were infected at a multiplicity of infection (MOI) of 5 with MERS-CoV and were incubated at 37°C for 1 h. After 1 hour, cells were aspirated, washed 1X with phosphate-buffered saline (PBS), and supplemented with fresh, pre-warmed Opti-MEM with 4% FetalClone II. Cells were then incubated at 37°C for an additional 19 hours.

SARS-CoV Urbani isolate (accession #MK062179) and SARS-CoV-2 USA-W1/2020 isolate (accession #MN985325) were gown in Vero E6 cells. Vero E6 cells were culture to ~90% confluence in T175 flasks. Immediately prior to infection the culture medium was aspirated and cells were washed with PBS. Flasks were infected at an MOI of 5 with SARS-CoV and MOI of 3 for SARS-CoV-2 at 37°C for 1 hour. After 1 hour cells were supplemented with pre-warmed DMEM with 5% FetalClone II. Cells were then incubated for an additional 24 hours at 37°C. To isolate virion RNA, cell supernatant was aspirated and concentrated using Amicon Ultra Centrifugal Filters (Millipore Sigma) to approximately 4 mL total volume. The supernatant was then lysed in TRIzol LS (Invitrogen), and viral RNA pellets were harvested according to the manufacturer’s suggested protocol. RNA was extracted from the TRIzol using chloroform followed by overnight precipitation at −20°C.

### SHAPE modification and library generation

Modified RNA was folded at 37°C in the presence of 10 mM MgCl2 and 111 mM KCl for 15 minutes, flowed by a 5-minute treatment of 100 nM 1-methyl-7-nitroisatoicanhydride (1M7) also at 37°C. Unmodified RNA was folded under the same conditions as the modified RNA and treated with DMSO for 5 minutes at 37°C. The denatured control RNA was incubated at 90°C for 2 minutes followed by a 2-minute treatment of 100 nM 1M7 at 95°C. Following all treatments, RNAs were purified using the RNA Clean & Concentrator −5 Kit (Zymo Research). Purified RNAs were random primed using Random Primer 9 (NEB) by incubation at 65°C for 5 minutes followed by rapid cooling on ice. They were then mixed with a master mix that consisted of 10 mM dNTPs, 0.1 M DTT, 500 mM Tris pH 8.0, 750 mM KCl, 500 mM MnCl_2_, and SuperScript II Reverse Transcriptase (Thermo Fisher Scientific). They were incubated at 25°C for 15 minutes, 42°C for 180 minutes, followed by a heat inactivation at 70°C for 15 minutes. After reverse transcription they were then cleaned using Illustra MicroSpin G-50 columns (GE Healthcare), double stranded DNA (dsDNA) was generated by use of NEBNext Ultra II Non-Directional RNA Second Strand Synthesis Module (NEB), purified using PureLink PCR Micro Kit (Thermo Fisher Scientific), and dsDNA was quantified using Qubit dsDNA HS Assay Kit (Thermo Fisher Scientific).

Libraries were prepared using Nextera XT DNA Library Preparation Kit (Illumina), cleaned using a 1:0.6 ratio of Agencourt AMPure XP (Beckman Coulter), and quantified again using Qubit dsDNA HS Assay Kit following manufacturer’s recommended protocol in each case. Libraries were sequenced on a MiSeq Desktop Sequencer (Illumina) using a MiSeq Reagent Kit v3 (600-cycle) (Illumina)^1, 2^.

### Structural prediction

Sequenced reads from the unmodified RNAs were aligned to the reference sequences downloaded from NCBI to correct viral sequences for potential mutations. SHAPE reactivities were derived using the ShapeMapper Pipeline v1.2. Structural predictions and Shannon entropy were obtained using Superfold v1.0 with a maximum pairing distance of 500 base pairs to calculate minimum free-energy models using SHAPE reactivities as folding constraints. Highly stable structured regions were predicted based on 55 base pair rolling averages of SHAPE reactivity (<0.3) and Shannon entropy (<0.04). Highly stable structured regions were expanded to encompass full base pairing regions as needed^3^. Specific structure’s minimum free energy models were generated using RNAstructure’s Fold v6.0.1 using standard settings and incorporating SHAPE reactivities as folding constraints^4^.

### Covariation and alignments

Homologous structures were found and base pairs with significant covariation were identified as in *Kutchko et al*^*1*^. Briefly, homologous structures were identified using the Infernal software suite v1.1.2^5^. A model was built for each structure using cmbuild and cmcalibrate. All available MERS-CoV (350 genomes), SARS-CoV (140 genomes), and SARS-CoV-2 (1,524 genomes) sequences available on ViPR on 5-12-2020 were searched for similar structures using cmsearch with -A option to automatically generate an alignment.

Base pairs with significant covariance were identified using R-scape program v1.2.3^6^.

MUSCLE alignments were performed on the CLC Genomics Workbench v12.0.3 (Qiagen) module Additional Alignments v1.9 for mutational frequency identification using the above mentioned ViPR sequences. Genomes containing gaps (>5 nucleotides) or multiple unidentified nucleotides were excluded from alignments. ClustalOmega alignments were performed using EMBL-EBI for individual viral genome comparisons^7^.

### Sequence conservation and structuredness

A sequence conservation score, ranging from 0 to 1, was calculated at each aligned genomic position after insertions and deletions were removed from the alignment. Whole genome conservation was calculated based on an average of each position’s conservation score across the entire genome and also the average of 67 base pair rolling windows across the entire genome. Structuredness was calculated at each nucleotide position by adding the SHAPE reactivity value to Shannon entropy value, both normalized to their mean value over the entire genome. This was used to create a ranking of each highly stable structured regions. Statistical analysis was performed using ANOVA: Single factor.

## Acknowledgements

We would like to thank the Moorman, Baric, and Heise labs for helpful conversations. This work was supported by the following grants from the National Institute of Allergy and Infectious Disease: AI137887 and AI138056 to N.J.M. and M.T.H., AI108197 to R.S.B., and support from T32 AI007419 to E.A.M. and E.J.F.

## Author Contributions

W.S., M.T.H., R.S.B., and N.J.M. conceptualized the work; W.S., E.J.F., and R.L.G. acquired data. W.S., E.A.M., and N.J.M. analyzed data; W.S., H.A.V., and N.J.M. drafted and revised the manuscript.

## Additional Information

Supplementary Information is available for this paper.

## References

1. Salata C, Calistri A, Parolin C, et al. Coronaviruses: a paradigm of new emerging zoonotic diseases. Pathog Dis 2019; 77 2020/02/18. DOI: 10.1093/femspd/ftaa006.

2. Hui DSC and Zumla A. Severe Acute Respiratory Syndrome: Historical, Epidemiologic, and Clinical Features. Infect Dis Clin North Am 2019; 33: 869–889. 2019/11/02. DOI: 10.1016/j.idc.2019.07.001.

3. Azhar EI, Hui DSC, Memish ZA, et al. The Middle East Respiratory Syndrome (MERS). Infect Dis Clin North Am 2019; 33: 891–905. 2019/11/02. DOI: 10.1016/j.idc.2019.08.001.

4. Tse LV, Meganck RM, Graham RL, et al. The Current and Future State of Vaccines, Antivirals and Gene Therapies Against Emerging Coronaviruses. Front Microbiol 2020; 11: 658. 2020/05/12. DOI: 10.3389/fmicb.2020.00658.

5. Yang D and Leibowitz JL. The structure and functions of coronavirus genomic 3’ and 5’ ends. Virus Res 2015; 206: 120–133. 2015/03/05. DOI: 10.1016/j.virusres.2015.02.025.

6. Sheahan TP, Sims AC, Zhou S, et al. An orally bioavailable broad-spectrum antiviral inhibits SARS-CoV-2 in human airway epithelial cell cultures and multiple coronaviruses in mice. Sci Transl Med 2020; 12 2020/04/08. DOI: 10.1126/scitranslmed.abb5883.

7. de Wit E, Feldmann F, Cronin J, et al. Prophylactic and therapeutic remdesivir (GS-5734) treatment in the rhesus macaque model of MERS-CoV infection. Proc Natl Acad Sci U S A 2020; 117: 6771–6776. 2020/02/15. DOI: 10.1073/pnas.1922083117.

8. Gorbalenya AE, Enjuanes L, Ziebuhr J, et al. Nidovirales: evolving the largest RNA virus genome. Virus Res 2006; 117: 17–37. 2006/03/01. DOI: 10.1016/j.virusres.2006.01.017.

9. Ganser LR, Kelly ML, Herschlag D, et al. The roles of structural dynamics in the cellular functions of RNAs. Nat Rev Mol Cell Biol 2019; 20: 474–489. 2019/06/12. DOI: 10.1038/s41580-019-0136-0.

10. Madhugiri R, Fricke M, Marz M, et al. Coronavirus cis-Acting RNA Elements. Adv Virus Res 2016; 96: 127–163. 2016/10/08. DOI: 10.1016/bs.aivir.2016.08.007.

11. Goebel SJ, Hsue B, Dombrowski TF, et al. Characterization of the RNA components of a putative molecular switch in the 3’ untranslated region of the murine coronavirus genome. J Virol 2004; 78: 669–682. 2003/12/25. DOI: 10.1128/jvi.78.2.669-682.2004.

12. Plant EP and Dinman JD. The role of programmed-1 ribosomal frameshifting in coronavirus propagation. Front Biosci 2008; 13: 4873–4881. 2008/05/30. DOI: 10.2741/3046.

13. Plant EP, Sims AC, Baric RS, et al. Altering SARS coronavirus frameshift efficiency affects genomic and subgenomic RNA production. Viruses 2013; 5: 279–294. 2013/01/22. DOI: 10.3390/v5010279.

14. Cantara WA, Olson ED and Forsyth KM. Progress and outlook in structural biology of large viral RNAs. Virus Res 2014; 193: 24–38. 2014/06/24. DOI: 10.1016/j.virusres.2014.06.007.

15. Siegfried NA, Busan S, Rice GM, et al. RNA motif discovery by SHAPE and mutational profiling (SHAPE-MaP). Nat Methods 2014; 11: 959–965. 2014/07/17. DOI: 10.1038/nmeth.3029.

16. Smola MJ, Rice GM, Busan S, et al. Selective 2’-hydroxyl acylation analyzed by primer extension and mutational profiling (SHAPE-MaP) for direct, versatile and accurate RNA structure analysis. Nat Protoc 2015; 10: 1643–1669. 2015/10/02. DOI: 10.1038/nprot.2015.103.

17. Yang D, Liu P, Wudeck EV, et al. SHAPE analysis of the RNA secondary structure of the Mouse Hepatitis Virus 5’ untranslated region and N-terminal nsp1 coding sequences. Virology 2015; 475: 15–27. 2014/12/03. DOI: 10.1016/j.virol.2014.11.001.

18. Chen SC and Olsthoorn RC. Group-specific structural features of the 5’-proximal sequences of coronavirus genomic RNAs. Virology 2010; 401: 29–41. 2010/03/06. DOI: 10.1016/j.virol.2010.02.007.

19. Di H, McIntyre AA and Brinton MA. New insights about the regulation of Nidovirus subgenomic mRNA synthesis. Virology 2018; 517: 38–43. 2018/02/25. DOI: 10.1016/j.virol.2018.01.026.

20. Sola I, Moreno JL, Zuniga S, et al. Role of nucleotides immediately flanking the transcription-regulating sequence core in coronavirus subgenomic mRNA synthesis. J Virol 2005; 79: 2506–2516. 2005/02/01. DOI: 10.1128/JVI.79.4.2506-2516.2005.

## Methods References

1. Kutchko KM, Madden EA, Morrison C, et al. Structural divergence creates new functional features in alphavirus genomes. Nucleic Acids Res 2018; 46: 3657–3670. 2018/01/24. DOI: 10.1093/nar/gky012.

2. Smola MJ, Rice GM, Busan S, et al. Selective 2’-hydroxyl acylation analyzed by primer extension and mutational profiling (SHAPE-MaP) for direct, versatile and accurate RNA structure analysis. Nat Protoc 2015; 10: 1643–1669. 2015/10/02. DOI: 10.1038/nprot.2015.103.

3. Siegfried NA, Busan S, Rice GM, et al. RNA motif discovery by SHAPE and mutational profiling (SHAPE-MaP). Nat Methods 2014; 11: 959–965. 2014/07/17. DOI: 10.1038/nmeth.3029.

4. Reuter JS and Mathews DH. RNAstructure: software for RNA secondary structure prediction and analysis. BMC Bioinformatics 2010; 11: 129. 2010/03/17. DOI: 10.1186/1471-2105-11-129.

5. Nawrocki EP and Eddy SR. Infernal 1.1: 100-fold faster RNA homology searches. Bioinformatics 2013; 29: 2933–2935. 2013/09/07. DOI: 10.1093/bioinformatics/btt509.

6. Rivas E, Clements J and Eddy SR. A statistical test for conserved RNA structure shows lack of evidence for structure in lncRNAs. Nat Methods 2017; 14: 45–48. 2016/11/08. DOI: 10.1038/nmeth.4066.

7. Madeira F, Park YM, Lee J, et al. The EMBL-EBI search and sequence analysis tools APIs in 2019. Nucleic Acids Res 2019; 47: W636–W641. 2019/04/13. DOI: 10.1093/nar/gkz268.

